# Zika virus hijacks extracellular vesicle tetraspanin pathways for cell-to-cell transmission

**DOI:** 10.1101/2021.03.02.433679

**Authors:** Sara B. York, Li Sun, Allaura S. Cone, Leanne C. Duke, Mujeeb R. Cheerathodi, David G. Meckes

## Abstract

Extracellular vesicles (EVs) are membrane-encapsulated structures released by cells which carry signaling factors, proteins and microRNAs that mediate intercellular communication. Accumulating evidence supports an important role of EVs in the progression of neurological conditions and both the spread and pathogenesis of infectious diseases. It has recently been demonstrated that EVs from Hepatitis C virus (HCV) infected individuals and cells contained replicative-competent viral RNA that was capable of infecting hepatocytes. Being a member of the same viral family, it is likely the Zika virus also hijacks EV pathways to package viral components and secrete vesicles that are infectious and potentially less immunogenic. As EVs have been shown to cross blood-brain and placental barriers, it is possible that Zika virus could usurp normal EV biology to gain access to the brain or developing fetus. Here, we demonstrate that Zika virus infected cells secrete distinct EV sub-populations with specific viral protein profiles and infectious genomes. Zika virus infection resulted in the enhanced production of EVs with varying sizes and density compared to those released from non-infected cells. We also show that the EV enriched tetraspanin CD63 regulates the release of EVs, and Zika viral genomes and capsids following infection. Overall, these findings provide evidence for an alternative means of Zika virus transmission and demonstrate the role of EV biogenesis and trafficking proteins in the modulation of Zika infection.

**Importance:** Zika virus is a re-emerging infectious disease that spread rapidly across the Caribbean and South America. Infection of pregnant women during the first trimester has been linked to microcephaly, a neurological condition where babies are born with smaller heads due to abnormal brain development. Babies born with microcephaly can develop convulsions and suffer disabilities as they age. Despite the significance of Zika virus, little is known about how the virus infects the fetus or causes disease. Extracellular vesicles (EVs) are membrane-encapsulated structures released by cells that are present in all biological fluids. EVs carry signaling factors, proteins and microRNAs that mediate intercellular communication. EVs have been shown to be a means by which some viruses can alter cellular environments and cross previously unpassable cellular barriers. Thus gaining a greater understanding of how Zika affects EV cargo may aid in the development of better diagnostics, targeted therapeutics and prophylactic treatments.

## INTRODUCTION

Zika virus (ZIKV) is a single-stranded, positive sense RNA virus (ss+RNA) belonging to the *Flaviviridae* family (1). This reemerging arbovirus is primarily transmitted to humans by *Aedes* species mosquitos, but additional evidence indicates that it can be transmitted sexually and vertically from mother to fetus (2, 3). Historically, ZIKV has been classified as a self-limiting febrile disease and until more recently, reported human cases have been sporadic and clustered in specific geographical regions. Since the 2015 outbreak in Brazil, the virus has continued to spread in the western hemisphere and the number of Zika infections surged with reports of neurological diseases, such as Guillain-Barré syndrome and congenital microcephaly associated with infection (4–6). The increased global threat of Zika congenital syndrome and Zika-associated neuropathies prompted the scientific community to gain a greater understanding of Zika virus replication and pathogenesis in the host. However, the important question of how this flavivirus is able to cross placental and blood brain barriers to infect neural progenitor cells of the developing fetus remains unanswered.

EVs are a heterogeneous mixture of communicative vehicles secreted by cells in both healthy and pathogenic circumstances. EVs are typically classified based on size as well as cellular origin and include apoptotic bodies, microvesicles, and exosomes. Apoptotic bodies are large vesicles shed by cells undergoing apoptosis whereas, microvesicles are considered to be between 100-1000 nm in size and formed by budding at the cell surface (7, 8). Exosomes range between 40-150 nm in size and are derived endocytically through budding events inside the multivesicular body (MVB) (8). In addition to these major classes of vesicles, various microvesicle and exosome sub-populations likely exist with distinct cargos and functions (9).

Many viruses, including some flaviviruses, have been found to highjack EV biogenesis pathways for virus assembly and egress, or the secretion of specific viral RNAs and proteins (10–14). More recently, EVs have been found to be a unique means of transmission of certain enveloped and even non-enveloped viruses distinct from virion-mediated infection (15–17). There is a striking similarity between non-infectious EVs and virions. EVs from non-infected cells also resemble EVs from infected cells and can contain viral proteins and genome fragments (18). Historically virologists have referred to these EVs as non-infectious or defective particles. The similar biophysical properties make true separation of virions from EVs very difficult. In this study we were interested in analyzing EV subpopulations for Zika viral proteins and RNA to determine the effect of infection on all EVs populations.

EVs arise from every cell type, are present in all bodily fluids (19, 20) and have even been found to cross the blood brain barrier (21, 22) and the placenta (23, 24). Pregnancy increases the circulation of fetal EVs in maternal blood as early as the first trimester (23), which also coincides with the highest reported risk of infecting the fetus with ZIKV (25). Since virally-modified EVs have been found to manipulate cellular microenvironments and enhance virus transmission and pathogenicity (26–28), it is conceivable that EVs released from Zika infected cells may similarly affect the host.

How exactly viruses influence the cargo and functions of EVs is largely unknown. Nonetheless, it is likely at least in part by interactions between viral proteins and host EV trafficking and biogenesis pathways. Multiple pathways have been described in the literature. For instance, the endosomal sorting complex required for transport (ESCRT) pathway is important for invagination of the MVB to form the intraluminal vesicles, which become exosomes when released from the cell (29, 30). Though, exosome biogenesis and release can also occur in an ESCRT- independent manner, as tetraspanins and the lipid ceramide have been shown to be important in exosome biogenesis and protein trafficking to the MVB (31, 32). Tetraspanins are a group of proteins containing four transmembrane domains, two extracellular loops, large and small, and three short intracellular regions (33). Tetraspanins are essential for many different cellular mechanisms, such as immune function (34) or trafficking of partner proteins to different organelles (35). Both ESCRT proteins and tetraspanin proteins are enriched in exosomes (36, 37).

There have been multiple established associations between viruses and EV biogenesis pathways. Human immunodeficiency virus (HIV), herpes simplex 1 (HSV1), Ebola virus, and rabies virus have been previously reported to hijack parts of the ESCRT pathway for assembly and egress (38–40). Additionally, human herpesvirus-6 (HHV-6) is believed to use the tetraspanin CD63 during replication (41). Other flaviviruses, such as dengue virus, were found to recruit several host ESCRT subunits to sites of replication and that these ESCRT subunits are required for efficient viral budding (42). Here, we sought to determine if Zika virus utilizes EV biogenesis pathways for virus replication and spread. Our overall hypothesis is that ZIKV usurps EV trafficking pathways in order to package viral genomic RNA and proteins to increase infectivity, evade the host immune response and possibly alter tropism of the virus.

## METHODS

### Cell culture

Vero E6 cells (a kind gift from Dr. Hengli Tang, Florida State University) were cultured in Dulbecco modified Eagle medium (Sigma, D5796) supplemented with 10% fetal bovine serum (Gibco, 26140-079), 2 mM L-glutamine (Corning; 25-005-CI), 100 IU of penicillin-streptomycin (Corning; 30-002-CI), and 100 μg/mL:0.25 μg/mL antibiotic/antimycotic (Corning; 30-002-CI). The SNB-19 cells were obtained from Charles River Laboratories, Inc. under contract of the Biological Testing Branch of the National Cancer Institute. SNB-19 cells were grown in RPMI 1640 (Sigma, R8758) supplemented with 10% fetal bovine serum (Gibco, 26140-079), 2 mM L-glutamine (Corning; 25-005-CI), 100 IU of penicillin-streptomycin (Corning; 30-002-CI), and 100 μg/mL:0.25 μg/mL antibiotic/antimycotic (Corning; 30-002-CI). MCF10a (ATCC, CRL-10317) cells were grown in MEBM (Lonza) with additives (MEGM kit, Lonza CC-3150) with 100 ng/mL cholera toxin (Sigma).

### Extracellular vesicle depleted FBS

EV depleted FBS was used in all of the experiments in which EVs were harvested. FBS was centrifuged at 100,000 rcf for 20 hours in a SW32Ti rotor and then filtered through a 0.2 um filter.

### Plasmid cloningplenti X1 shRNA plasmids

CD9_shRNA_fwd

5′GATCCC CAAGAAGGACGTACTCGAAAC GTGTGCTGTCC GTTTCGAGTACGTCCTTCTTG TTTTTGGAAA

CD9_shRNA_rvs

5′AGCTTTTCCAAAAA CAAGAAGGACGTACTCGAAAC GGACAGCACAC GTTTCGAGTACGTCCTTCTTG GG

CD63 shRNA_ fwd

5′GATCCCCAACGAGAAGGCGATCCATAAGTGTGCTGTCCTTATGGATCGCCTTCTCGTTGTCTTTTTGGAAA

CD63 shRNA_ rvs

5′AGCTTTTCCAAAAACAACGAGAAGGCGATCCATAAGGACAGCACACTTATGGATCGCCTTCTCGTTGGG

CD81 shRNA_fwd

5’GATCCCACATCCTGACTCCGTCATTTA GTGTGCTGTCC<u>TAAATGACGGAGTCAGGATGT</u>TTTTTGGAAA

CD81 shRNA_rvs 5’

AGCTTTTCCAAAAAACATCCTGACTCCGTCATTTAGGACAGCACTAAATGACGGAGTCAGGATGTGG

### Retrovirus production

Transduction retrovirus particles were collected from HEK293T cells following Lipofectamine 3000 transfection of expression plasmids (pLenti CMV TetR BLAST and plenti X1 shRNA plasmids) and packaging plasmids pMD2.G (Addgene; number 12259; a gift from Didier Trono) and PSPAX2 (Addgene; number 12260; a gift from Didier Trono) according to the manufacturer’s instructions (Invitrogen, L3000015). The overexpression pCT-CD9-RFP (SBI, CYTO123-PA-1) and pCT-CD63-GFP (SBI, CYTO120-PA-1) plasmids were transfected in HEK293T cells with the packaging plasmids pMD2.G (Addgene; number 12259; a gift from Didier Trono), pMDLgpRRE (Addgene; number 12251; a gift from Didier Trono), pRSVRev (Addgene; 12253; a gift from Didier Trono) according to the manufacturer’s instructions (Invitrogen, L3000015). Medium was collected and reapplied at 48, 72, and 96 h following transfection, centrifuged for 10 min at 1,000 rcf, filtered through a 0.45 μm filter, and frozen at −80°C until use.

### Generation of cell lines

Cells stably expressing shRNA of different genes under the control of a tetracycline-inducible promoter were created by first transducing SNB-19 cells with lentivirus particles containing pLenti CMV TetR BLAST (Addgene; number 17492). The cells underwent selection with media containing 10 μg/mL of blasticidin (Invivogen; ant-bl-1) for two weeks. These cells were then transduced with plenti X1/zeo shCD63, plenti X1/puro shCD9, or plenti X1/zeo shCD81. Doubly stable cells were selected with medium supplemented with blasticidin (10 μg/mL) and puromycin (2 μg/mL) or zeocin (100 μg/mL concentration) for 2 weeks. The inducible shRNA cell lines were tested for knockdown by immunoblotting of cell lysates following 24 hours of doxycycline induction. SNB-19 cells were transduced with lentiviral particles containing CD9-RFP (SBI, CYTO123-PA-1) or CD63-GFP (SBI, CYTO120-PA-1) and selected with media containing puromycin (2 μg/mL) for two weeks. SNB-19 cells were transfected (lipofectamine 3000; Invitrogen, L3000015) with the overexpression mCherry CD81 (gift from Michael Davidson, Addgene plasmid #55012) construct and selected with media containing neomycin (500 μg/mL; Corning; 30-234-CI) for two weeks. Overexpression cell lines were confirmed by immunoblotting of whole cell lysate and live cell imaging.

### Zika virus stock preparation

All experiments were conducted using the Zika PRVABC-59 strain (kind gift from Dr. Hengli Tang, Florida State University). Viral stocks were grown in Vero E6 cells (kind gift from Dr. Hengli Tang, Florida State University) and plaque assays were used to determine virus stock titers. The protocol for producing, harvesting and enumerating Zika virus stocks was followed as described in Agbulos et al., 2016 (43).

### Zika infection

Cells were counted with an automated cell counter (Cellometer Vision, software version 2.1.4.2; Nexcelom Biosciences) prior to seeding. Cells were seeded in normal growth medium for cell type with 10% FBS. For the shRNA inducible stables, cells were induced with 1 μg/mL doxycycline 16-18 hours prior to infection. After 16 -18 hours, the media was removed and cells were infected with Zika virus (MOI=0.01-0.05) or mock (just media) for one hour. The flasks were rocked every 10 minutes. The medium was removed, the flasks were washed with phosphate-buffered saline (PBS), and growth media was added (1 μg/mL doxycycline was reapplied to shRNA cells). After 24 hours, the medium was removed and media containing 10% vesicle-depleted FBS (1 μg/mL doxycycline was added to shRNA cells) was applied. Following 48-72 hours, the media and cells were harvested for further processing.

### Extracellular vesicle enrichment

Extracellular vesicles were isolated and characterized from cell-conditioned medium following the 2018 MISEV guidelines and as previously described (44–46). Media was collected and centrifuged serially (500 rcf for 5 minutes, 2,000 rcf for 10 minutes, and 10,000 rcf for 30 minutes) to remove cell debris, apoptotic bodies and microvesicles. Supernatant was mixed 1:1 with a 16% PEG 6000 solution (16%, wt/vol, polyethylene glycol, 1 M NaCl) for a final concentration of 8% and incubated overnight at 4°C. The media/PEG solutions were centrifuged at 3,214 g for 1 hour the following day in order to obtain crude EV pellets. These pellets were resuspended in phosphate-buffered saline (PBS) and ultracentrifuged at 100,000 rcf for 2 hours. The EV pellets were resuspended in particle-free PBS (or sucrose buffer if to be further purified on a gradient) and stored at 4°C until additional analyses.

### Iodixinol density gradient EV subpopulation separation

For further purification and sub-population separation by floatation density gradient, EV pellets were instead resuspended in 1.5 mL of 0.25 M sucrose buffer (10 mM Tris, pH 7.4). Gradients (10-30%) were constructed as previously described in detail (9, 12) using OptiPrep (Sigma, D1556). EVs in the 1.5 mL were mixed 1:1 with 60% iodixanol to make a final concentration of 30% and loaded on the bottom of the gradient. Then, 1.3 mLs of 20% and 1.2 mLs of 10% were layered on top. The gradients were centrifuged for 1 hour at 50,000 rpm in a MLS 50 rotor (15min acceleration and de-acceleration for 90 min total time). The gradients were fractionated by pipetting 490 mL from the top. Ten 490 ul fractions were collected and the gradient pellet was resuspended as the 11^th^ fraction. Following fractionation, samples were washed in PBS and pelleted again by ultracentrifugation at 100,000 g for 2 hours. Pellets were resuspended in particle-free PBS.

### Immunoblot analysis

Whole-cell lysates were harvested by trypsinizing with 0.25% trypsin for 10 minutes, inactivating with growth media and pelleting at 500 × g for 5 min. Cells were lysed with radioimmunoprecipitation assay (RIPA) buffer as described previously (12). EVs were lysed using a strong lysis buffer containing urea (5% SDS, 10 mM EDTA, 120 mM Tris-HCl [pH 6.8], 8 M urea, protease inhibitor cocktail: Thermo; 78438). Cell lysate protein quantification was measured by Pierce 660 nm Protein Assay (Invitrogen, 22662) and the EZQ^TM^ Protein Quantitation kit (Thermo; R33200) method was used to quantify the EV lysates. Laemmli 5x sample buffer (with or without 2% BME for reducing or nonreducing conditions) was added to cell and EV lysates before SDS-PAGE separation. Equal protein of cell or EV lysate (or volume of EVs) was loaded into SDS polyacrylamide gels. Western blot analysis was performed as described previously (47). Ponceau S stain applied to visualize the total protein. Blots were probed using the following antibodies: Alix ([Q-19] Santa Cruz; SC-49268), HSC70 ([B-6] Santa Cruz; SC-7298), CD63 ([TS63] Abcam; ab59479), CD9 ([MM2/57] Milipore; CBL162), CD81 ([B399 1.3.3.22] GeneTex; GTX34568), calnexin ([H-70] Santa Cruz; sc-11397), Zika Envelope (EastCoast Bio; HM325), Zika capsid (GeneTex; GTX133317), Zika prM (GeneTex;133305), rabbit anti-mouse IgG (GeneTex; 26728), rabbit anti-goat IgG (GeneTex; 26741), or goat anti-rabbit IgG (Fab fragment) (GeneTex; 27171). Blots were imaged using an Image Quant LAS4000 (General Electric) and processed with ImageQuant TL v8.1.0.0 software, Adobe Photoshop CS6 and CorelDraw Graphic Suite X5.

### Nanoparticle tracking analysis

The Malvern NanoSight LM10 instrument was used for the nanoparticle tracking and the videos were processed using NTA 3.1 software with the camera level of 13 and a detection threshold of 3. Samples were diluted to be in a range of 2e8 to 1.8e9 particles and video length per replicate was 1 min.

### Electron microscopy

Electron microscopy grids were prepared and stained as previously described (47, 48). Grids (400 Hex Mesh Copper; Electron Microscopy Sciences, EMS) were exposed to vesicle preps contained in PBS for 1 hour. Following the incubation, the grids were washed with PBS three times before being fixed with 2% EM grade paraformaldehyde (EMS, EM Grade) for 10 minutes. The grids were washed again and incubated on a 2.5% glutaraldehyde solution (EMS, EM Grade) for another 10 mins. Then, the grids were washed 3 times with ultra-filtered water and stained with 2% uranyl acetate EMS). The grids were applied to 0.4% uranyl acetate/ 0.13% methylcellulose for 10 minutes and allowed to harden at room temperature for 24 hours. The imaging was performed on a CM Biotwin electron microscope.

### RNA isolation and Reverse transcription

Total RNA of cell or fraction samples were isolated by Trizol reagent (Invitrogen) and quantified by nanodrop. Less than 1ug of total RNA was used for reverse transcription by qScript cDNA SuperMix (Quantabio, 95048).

### Quantitative real-time PCR and Data analysis

RT-qPCR was preformed following the protocol from Xu et al. with a small modification (49). Standard 3-step cycles protocol (40 cycles of 95 ◦C for 5 s, 60 ◦C for 10 s, 72 ◦C for 20 s) was used instead of one-step fast 2-step cycles as the input was cDNA not RNA. Another set of primers targeting inner segment of Zika RNA were designed to confirm the whole genome packed in fractions. PerfeCTa SYBR® Green FastMix (Quantabio, 95072), assay primers and cDNA were prepared in 20 uL reaction and run on CFX96 qPCR machine (Bio-Rad). Zika RNA levels were measured by RT-qPCR and either normalized to GAPDH or calculated to copy number from an external standard with ΔΔCt method. A Zika standard curve was established to determine genome copies and the Bioanalyzer Agilent 2100 was utilized to confirm correct end product size.

### Primers sequence

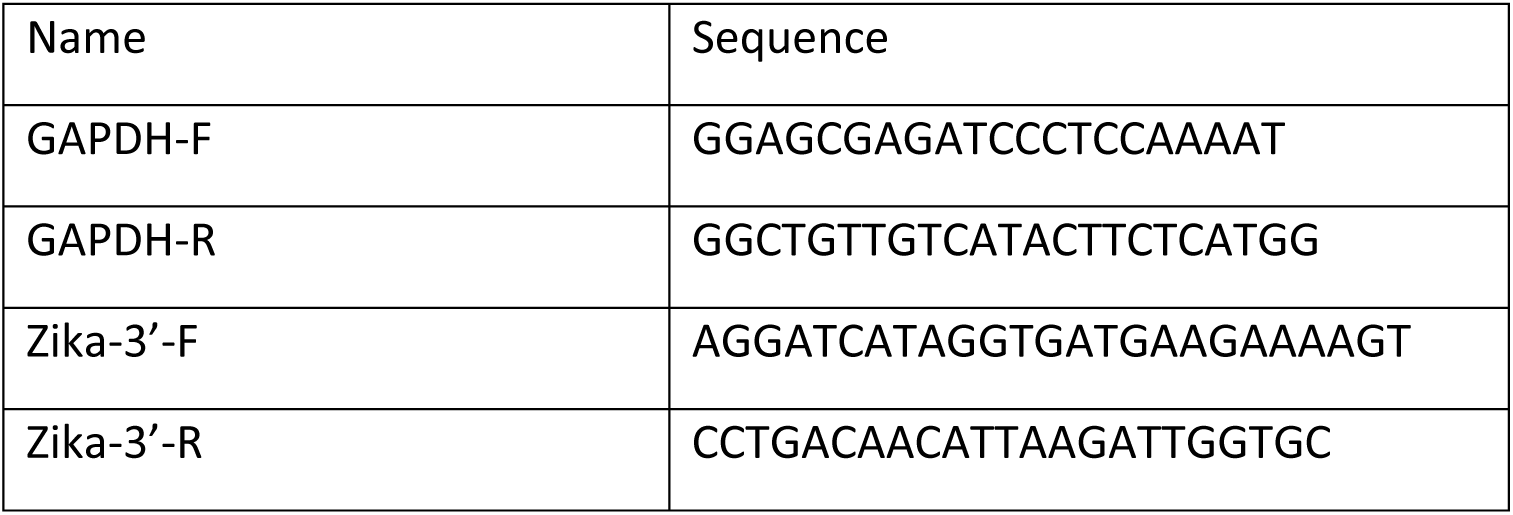
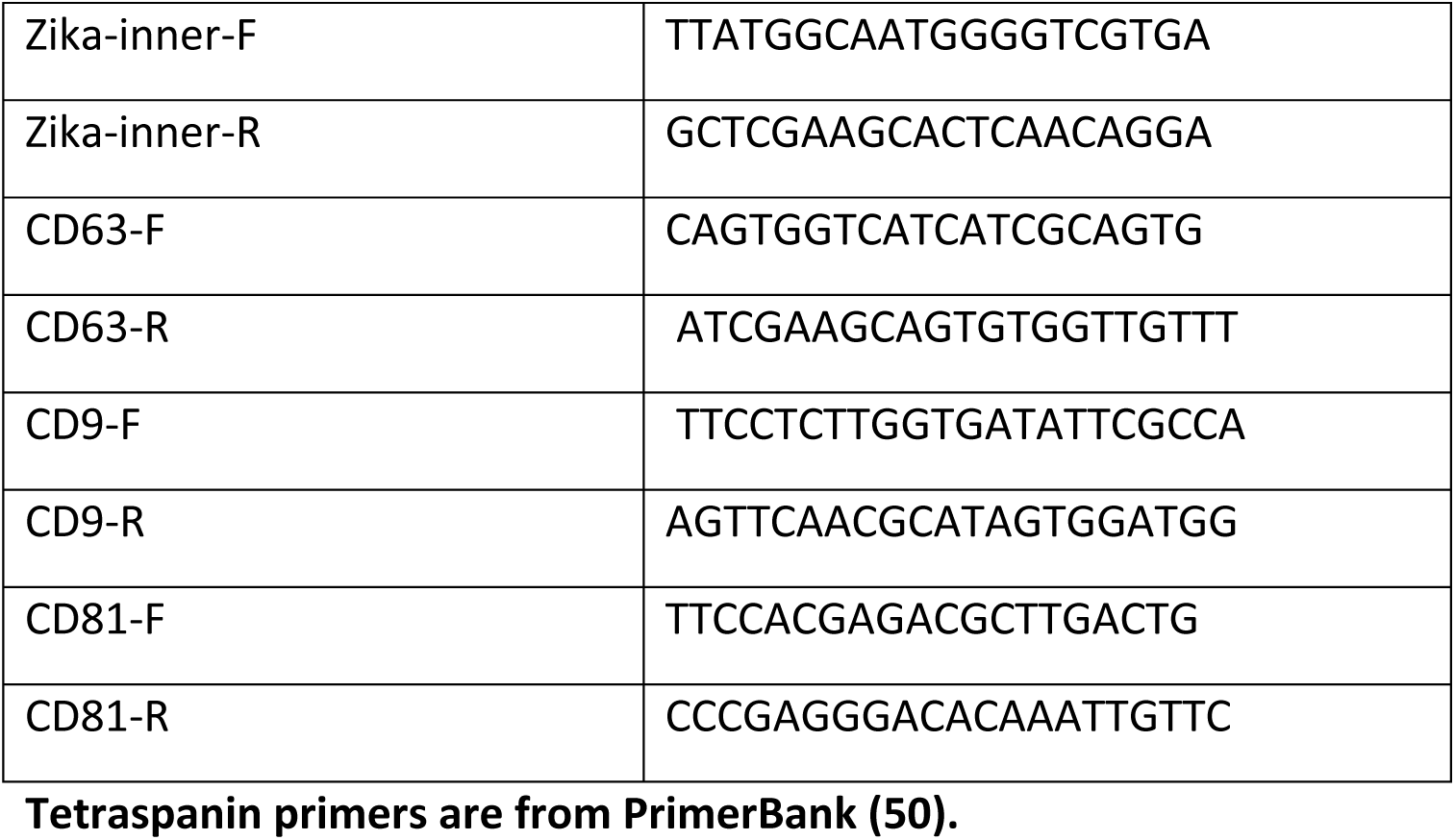

### Triton EV experiment

EVs were treated with RNase A (1mU) either with or without TritonX-100 (1%) for 15 mins. After adding RNase inhibitor, total RNAs were isolated by Trizol and Zika RNA were quantified by qPCR as described.

### EV Immunoprecipitation

Extracellular vesicles were harvested from the 100K pellet from Zika infected SNB-19 cells as previously described. Pre-conjugated anti-CD9 (Invitrogen #10614D), CD63 (Invitrogen #10606D), and CD81 (Invitrogen #10616D) antibodies were employed to pull-down EVs from the 100K Zika infected pellet. The MagCapture Exosome Isolation Kit PS (FujiFilm #293-77601) was utilized for the PS IP experiments. All IP experiments were performed as recommended by the manufacturer. Isolated EVs were further processed by WB, qPCR, and infectivity as previously described.

### EV transfer experiments

Total EVs were obtained by harvesting media from Zika infected SNB19 (control, induced shRNA and overexpression) cells or mock 72 hours post infection. The media was centrifuged for 1000 rcf for 10 minutes and filtered through a 0.45 μm filter. Then, the media was mixed 1:1 with 16% PEG and stored at 4°C overnight. The next day, the media was centrifuged at 3,214 rcf for 1 hour and total EVs were resuspended in sterile PBS.

EVs harvested from density gradients or total EVs were applied to SNB-19 cells seeded in 6 well plates. The cells were monitored every 24 hours for 72-96 hours and images were recorded using live cell imaging on the Keyence BZ-X700/BZ-X710 microscope. Cells were harvested and lysed, as previously described, 72-96 hours post infection. Cell lysate protein was quantitated before being separated on a SDS-PAGE gel and immunoblotted for Zika proteins.

### Statistical methods and analysis software

The Student’s two-sample *t* test or one-way ANOVA was used to determine significance. Prism8, Microsoft Excel, Adobe Photoshop CS6, and CorelDraw X5 were used in the design of the figures.

### Preparation of fixed cells for confocal microscopy

Infection: SNB-19 WT, inducible shRNA and overexpression cells were seeded on glass coverslips the day prior to infection with Zika virus. The following morning, shRNA cells were induced with 1 µg/mL doxycycline and 4-6 hours later all cell lines were infected with Zika virus at a MOI of 2. Following one-hour viral incubation at 37°C, media was replaced and doxycycline was reapplied to the shRNA cells. Coverslips were processed for IF and live cells were imaged 48 hours post infection.

Immunofluorescence (IF): Media was removed from wells containing the coverslips and the cells were fixed with 4% paraformaldehyde for 10 minutes. The coverslips were washed with PBS following fixation and permeabilized for 30 minutes with 0.2% Triton X-100 PBS (PBS-T). Cells were blocked in 5% goat serum in PBS-T for 30-60 minutes at room temperature. Primary antibody was added to coverslips (Envelope 1:500; Capsid 1:200; CD63 1:100 diluted in 5% goat serum/0.2%Tween PBS) and incubated at 4°C overnight. Primary antibody was removed and the coverslips were washed with PBS-T. Secondary anti-mouse (594, Thermo 35511; 488, Thermo 35503; 405, Rockland 610-146-002-0.5) or anti-rabbit (488, ab150077) at 1:1000 in 5% goat serum/PBS-T was applied and incubated at room temperature for 1 hour. The coverslips were washed again with PBS-T and mounted or stained with DAPI (Thermo, 6648) diluted 1:10,000 in PBS for 10 minutes. Coverslips were applied to mounting media (4% propyl gallate, 90% glycerol, PBS) on glass slides and allowed to harden overnight. Slides were imaged with a Zeiss LSM 880 microscope and images were processed using Zen 2.1 Black software.

## RESULTS

### Zika virus specifically modifies small EV density, cargo and secretion

In order to investigate whether Zika virus alters small EV populations, EVs were isolated by differential centrifugation with PEG precipitation and further purified the small EVs with a slightly modified iodixanol gradient method originally described by Kowal and colleagues (9, 51). For the gradient, EVs are loaded on the bottom of ultracentrifuge tubes and allowed to migrate up (i.e., float) into the gradient to their appropriate densities (9, 51). The paper by Kowal and colleagues, reported that small EVs travel to fraction 3 and 5, corresponding to approximately 1.11 and 1.15 g/mL respectively (9, 51). The EVs in fraction 3, termed small light EVs, were found to represent an enrichment in bon-a-fide exosomes of endosomal origin. The densities of the fractions from our gradient were measured and confirmed to yield similar densities as to those previously described in Kowal et al., 2016 (Fig 1A).

**Fig. 1.**
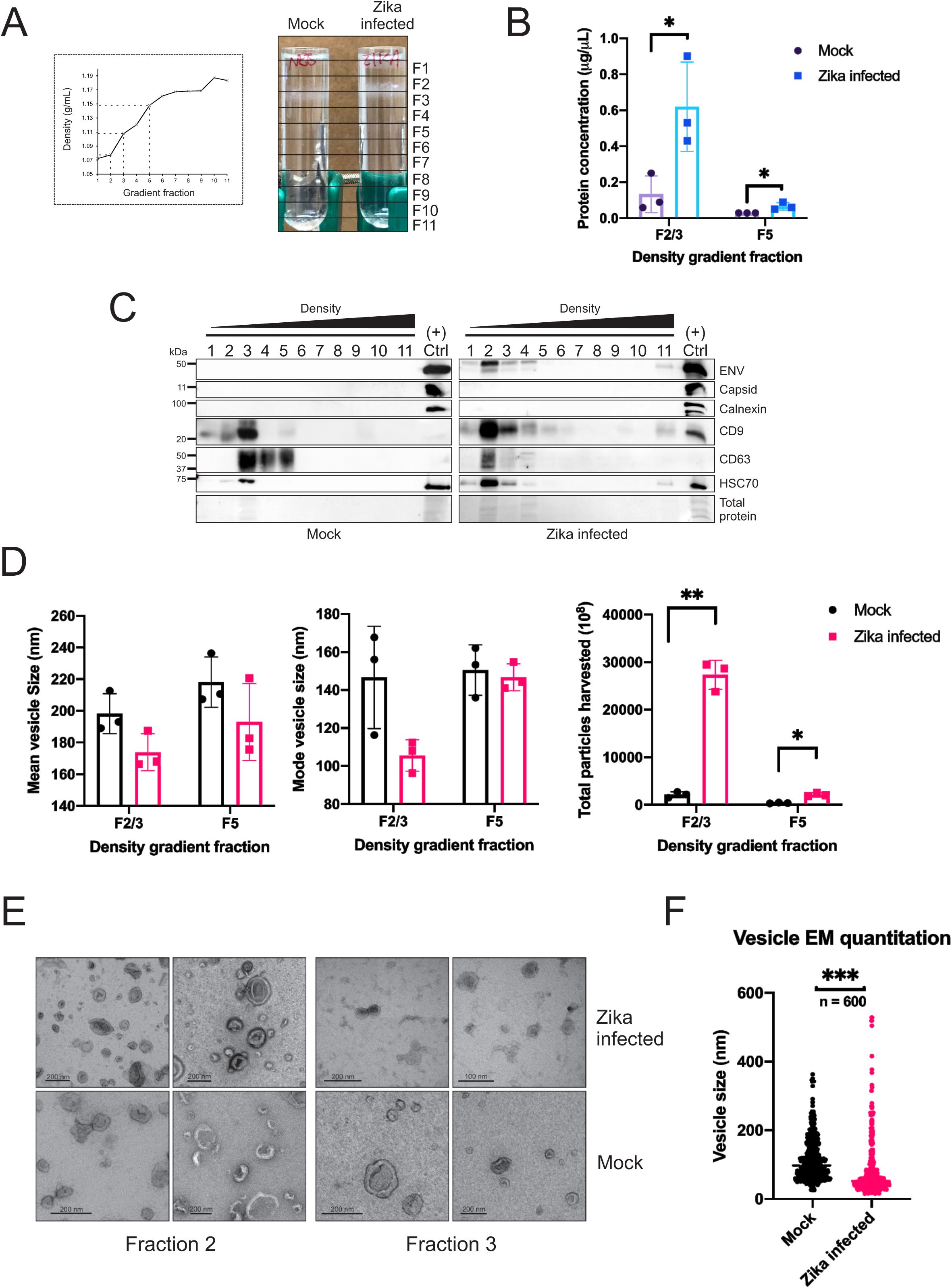
Zika virus specifically modifies small EV density, cargo and secretion. EV-depleted media (50-100 mL) harvested from SNB-19 glioblastoma (36-72e6) cells infected with Zika virus (MOI = 0.05-0.1) or mock for 72 hours, underwent differential centrifugation and PEG precipitation. The 100K EV pellet was further purified in a density gradient. A) Densities of the fractions were measured with a refractometer and visual differences between the uninfected and Zika-infected gradients was captured. B) Protein quantitation of fractions 2/3 and 5 from Zika-infected and uninfected cells (p<0.05). C) Equal volumes of the uninfected and Zika infected density fractions were separated by SDS-PAGE and immunoblotted for Zika proteins, tetraspanins, and EV markers. Positive control is the total vesicles from Zika infected cells following 1000 x g spin for 10 min and PEG precipitation. D) The size and numbers of vesicles from combined fractions 2/3 and 5, were further examined by nanoparticle tracking analysis (NTA) (* p<0.05; *** p<0.001). E) Electron microscopy representative images of the vesicle containing fractions from uninfected versus Zika-infected cells.) F) Quantification of vesicle sizes of the density fraction 2/3 EVs from the EM images (n = 600; p<0.001). All error bars depict standard deviation of at least three independent experiments.

Visual differences in the EV band size and location were immediately evident (1A). Protein quantitation of the combined fraction 2&3 and fraction 5 demonstrated that the total protein level is significantly higher in the fraction EVs from the Zika infected, corroborating the presence of more EV material (Fig. 1B). Immunoblotting confirmed that EV markers and Zika envelope were primarily present in fraction 2 as opposed to fraction 3 in the mock (Fig. 1C). The gradient purification also revealed that Zika capsid protein failed to be detected in the fractions (Fig 1C). The NTA and EM images showed that Zika virus increases the secretion of significantly smaller sized EVs (Fig. 1D-F). Altogether these data indicate that Zika virus is varying the secretion and cargo of small light EVs consistent with the features and markers of exosomes (9).

### Zika modified small EVs contain viral RNA and are infectious

Since we established that Zika virus is modifying EVs, we wanted to then assess whether these vesicles contain Zika genomic RNA. For this, a RT-qPCR method published by Xu et al. in 2016, was modified marginally and utilized to quantify Zika RNA genome copies in the gradient fractions (Fig 2A) (52). Two unique sets of primers were used to detect the presence of viral RNA genomes and confirmed the correct size of the end products with a Bioanalyzer (Fig 2A). Using RT-qPCR, we were able to demonstrate that all of the top 5 fractions contained Zika genomes, but the copy number was greatest in fraction 2 (Fig. 2B). In addition, the data show that Zika infection can be transmitted by EVs contained in this fraction by monitoring the cytopathic effects (CPE) and immunoblotting for Zika capsid protein in the cells exposed to these vesicles (Fig. 2C-D). While we would expect that EVs containing positive sense Zika genome to be infectious regardless of how it enters a cell, we cannot exclude the possibility that some virions contained in the gradient fractions could also produce productive infection. However, since no capsid protein was detected in the fractions and EV proteins were enriched, we anticipate this to be negligible.

**Fig. 2.**
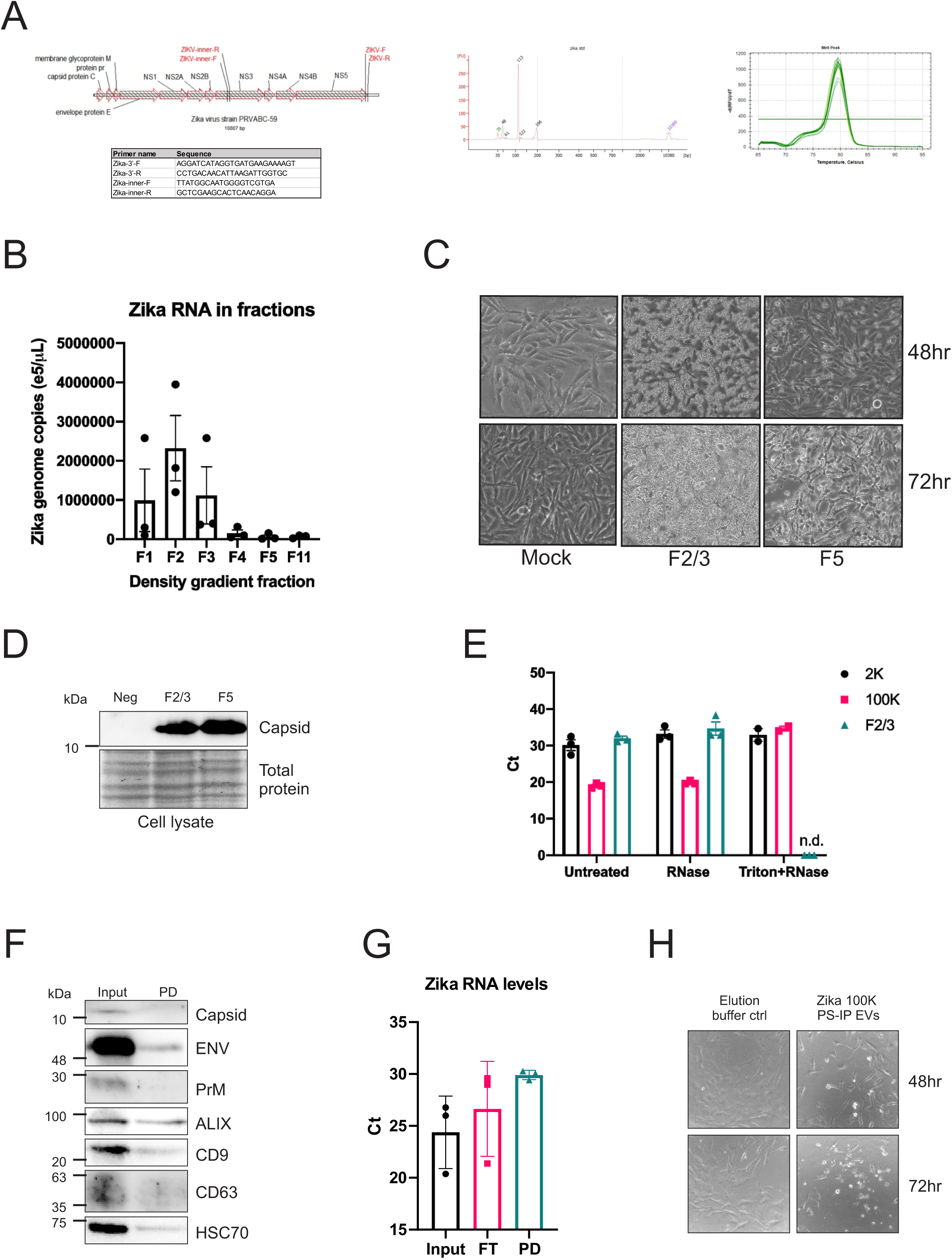
Zika modified small EVs contain viral RNA and are infectious. A) Previously published and newly designed primers were utilized for the detection of Zika virus genomes and full-length genomes. B) Zika genome copy number from gradient fractions (RT-qPCR of 3 independent experiments; error bars depict standard deviation). C) Equal volume of gradient fractions 2/3 and 5 were applied to uninfected SNB-19 cells. Cytopathic effects (CPE) were monitored and documented every 24 hours via live cell microscopy. D) Cells from the transfer experiment were harvested (72 hours post-infection), lysed and proteins were separated by SDS-PAGE. Immunoblot of cell lysates for Zika capsid protein to confirm infection. E) RT-qPCR results of gradient fraction 2/3, 100K and 2K untreated, treated with RNase, and with Triton and RNase. F) Immunoblot of EVs markers and Zika proteins of the input versus PS pull-downs of 100K Zika EVs. G) Zika RT-qPCR results of 3 independent input, flow through and PS 100K EV pull down. H) Equal volume of elution buffer only or PS pull down 100K EVs were added to Vero cells and CPE was captured by microscopy.

RNA can be found extracellularly in a non-membrane encapsulated form so, we wanted to evaluate whether the Zika RNA is actually contained within the vesicles or simply associated with the outside of the membrane. In order to assess this, RNA was extracted from Zika fraction 2/3, 100k and 2K EVs that were untreated, treated with RNase or treated Triton and RNase. The 2K EV were obtained from the pellet following a 2000xg spin and likely include large EVs as well as capsid containing virions. The 100K pellet is obtained following the differential centrifugation but prior to gradient purification and is expected to contain both capsid-containing virions and capsid-less Zika-modified EVs. Treating the EVs with RNase resulted in no significant increase in Ct in any of the EV groups, whereas with the addition of Triton prior to RNase treatment resulted in a lack of RNA detection in the fraction EVs (Fig. 2E). As would be expected of encapsulated genomes, there was also a detergent and Triton resistant genome population in the 100K fraction indicating the presence of virions in this pellet (Fig. 3E). These results support Zika RNA being contained within vesicles in the fraction 2/3 EVs since the RNA was not degraded by RNase treatment unless the membrane was first solubilized with detergent. The inability to detect any RNA in the Triton treated fraction 2/3 sample also supports that few if any genomes are present within a mature capsid found in virions.

**Fig. 3.**
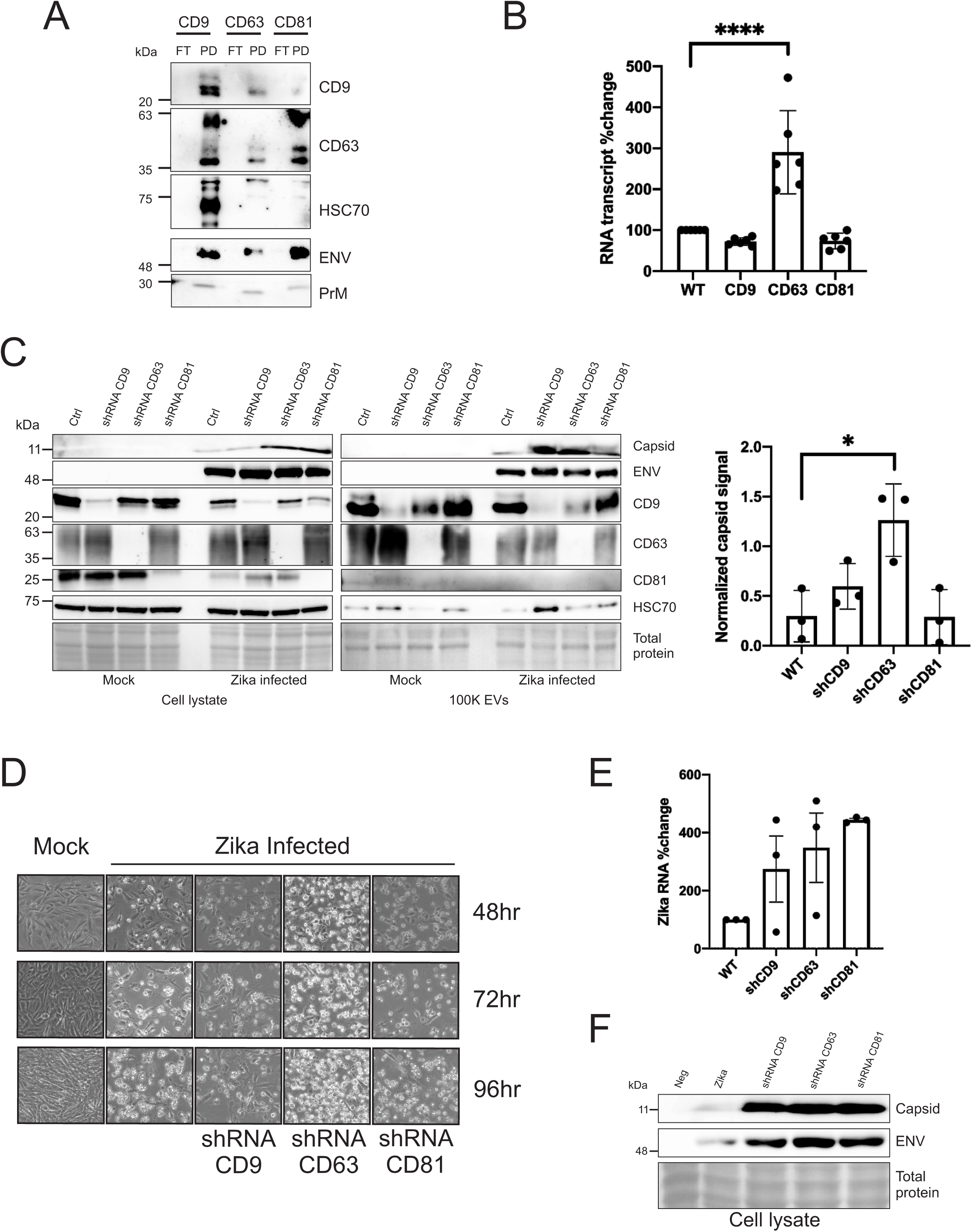
Knockdown of tetraspanins increases Zika infectivity and secretion of infectious particles/vesicles. A) Immunoblot of tetraspanin CD9, CD63, and CD81 EV immunoprecipitated proteins probed for EV markers and Zika proteins. B) RNA was extracted from cells 48 hours post Zika infection and RT-qPCR was performed to obtain transcript levels of CD9, CD63, CD81 (error bars represent SEM of 3 independent experiments; samples normalized to control and control set to 100). C) Immunoblots of cell and EV 100K lysate harvested 72 hours from uninfected or Zika infected ctrl and inducible tetraspanin knockdown cells that were probed for EV markers and Zika proteins. Quantitation of 100K EV Zika capsid levels of 3 independent experiments (error bars displayed as SEM; *p<0.05) D) Tetraspanin inducible shRNA knockdowns and control cells were monitored for CPE via live cell imaging every 24 hours for 4 days post-infection with Zika or mock. E) Quantitation of the percent change in Zika 100K EV RNA levels of 3 independent experiments (error bars display SEM of 3 independent experiments; samples normalized to control and control set to 100). F) Immunoblot of cell lysate from cells exposed to EVs from tetraspanin knockdown cells.

To provide more evidence that these EVs represent a distinct infectious population, we then decided to perform an alternative isolation method utilizing beads conjugated with a phosphatidylserine (PS) binding protein to pull down EVs because EVs are known to have exposed PS on their surface (53, 54). The 100k pellet was utilized for these experiments since it is expected to contain both virions and Zika-modified EVs. Immunoblotting of the PS pull-down EV proteins revealed that EV markers (Alix, CD63, CD9, and HSC70) and Zika envelope were captured but the Zika capsid and PrM proteins were only detected in the input (Fig. 3F). qPCR was then performed on these EVs and found that Zika RNAs were detected in both the pull-down and flow through suggesting that this pellet likely contains both virions and Zika-modified EVs (Fig. 3G). We analyzed the infectivity of the EV pull downs by introducing them to naïve Vero cells following elution from the beads and monitoring CPE over 72 hours (Fig. 3H). CPE was visualized in the Vero cells exposed to the Zika PS pulled-down EVs but not in the buffer control indicating that these EVs are indeed infectious (Fig. 3H). Altogether, these data provide evidence for Zika virus modifying EV cargo to produce infectious EV subpopulations distinct from virions.

### Knockdown of tetraspanins increases Zika infectivity and secretion of infectious particles/vesicles

To further characterize the subpopulations of Zika-modified EVs, immunoprecipitation of the 100K pellet was performed with antibody conjugated beads to CD9, CD63, and CD81 since tetraspanins are known EV markers that have significant roles in EV biogenesis. Western blotting of the IP proteins demonstrated that both EV markers (CD9, CD63 and HSC70) and Zika proteins (Envelope and PrM) were captured in all three pull-downs and not detected in the flow through (Fig. 3A). These data suggest that Zika virions may contain tetraspanins in their membranes since little protein was detected in the flow-through. However, this may not be surprising since prior studies have shown that some enveloped viruses bud from tetraspanin-enriched microdomains and thus contain tetraspanins on the surface (58, 59).

EV trafficking and biogenesis proteins, including tetraspanins, have been previously shown to have a role in the packaging of viral proteins into EVs and virus assembly and egress (12, 55). As Zika appeared to be enhancing the production of EVs from infected cells with infectious genomes, we reasoned that the virus may be influencing expression of these genes. Therefore, we assessed whether Zika infection altered the transcript levels of tetraspanin genes in cells. The 48-hour time point was chosen as substantial CPE usually occurs at 72 hours and the levels should be evaluated while the cells were infected but before extensive cell death. The results show that CD63 transcript levels increase considerably but CD9 and CD81 levels decrease slightly (Fig. 3B). Since these transcripts were measured at 48 hours during early infection, it is possible the virus requires CD63 during a stage of the replication cycle. The elevated CD63 transcripts could also represent a host reaction to the viral infection rather than an intended outcome of the virus.

To test the importance of tetraspanins in Zika replication, a doxycycline inducible shRNA system was employed to knockdown the tetraspanin proteins CD9, CD63 and CD81 in infected cells. The tetraspanin knockdown cell lines were infected with a MOI of 0.5. No differences in CPE or Zika protein levels were detected between the infected SNB-19 WT and inducible shRNA scramble cells (data not shown). Immunoblots of the cell and EV lysate demonstrate efficient knockdowns of the tetraspanins (Fig. 3C). In addition, there were increased capsid levels in the CD9 and CD63 knockdowns, but a statistically significant increase was only observed for the EV shCD63 lysate (Fig. 3C). Although alterations in capsid protein levels were quantifiable, Zika envelope levels appeared relatively unaffected (Fig. 3C). The CPE was monitored by live cell imaging for 4 days and substantial CPE was visualized in the CD63 knockdown cell line when compared with the other cell lines in all three time points (Fig. 3D). RT-qPCR of the EVs revealed a considerable increase in the Zika RNA from all three of the tetraspanin shRNA cells (Fig. 3E). Therefore, tetraspanin, especially CD63, appear to modulate EV, capsid, and genome release from infected cells.

Next, we wanted to analyze the infectious capability of the viral particles/vesicles. This was accomplished by transferring tetraspanin knockdown total EVs to naïve SNB-19 cells and harvesting the cells 96 hours post-exposure. Immunoblots of the knockdown exposed cells reveal dramatic increases in Zika capsid protein levels but the envelope protein was only increased moderately (Fig. 3F). Altogether these data suggest that EV tetraspanins CD9, CD63, and CD81 modulate Zika replication and spread.

### Overexpression of tetraspanin levels decreases Zika infectivity and transmission

Since knocking down tetraspanins seems to increase EV and genome release, we then wanted to explore the effects of tetraspanin overexpression. When SNB-19 WT cells and SNB-19-GFP cells were infected, no difference was observed in the resulting CPE or cellular Zika protein levels confirming that overexpression of GFP alone does not influence viral infection (data not shown). Zika infected SNB-19 cells overexpressing CD9, CD63, and CD81 and total EV lysates (2K, 10K, and 100K vesicles and viral particles) were then harvested following 96 hours post-infection and separated on SDS-PAGE gels. Cell lysate blots demonstrate an effective overexpression of tetraspanins in all three cell lines (Fig. 4A). Zika capsid and envelope levels detected in the cell lysates of the CD63 overexpression were decreased when compared with the control and other cell lines (Fig. 4A). Immunoblot of total EVs revealed a substantial decrease in capsid levels from the CD9 and CD63 overexpression cells while envelope protein was only mildly decreased (Fig. 4B). This was a striking finding since the total EV prep should include Zika virions and Zika modified EVs. This implies that overexpression of CD9 and CD63 disrupted the secretion of Zika virions since capsid is a critical structural component of the virion (56). The total EVs of all three overexpression cells were analyzed by RT-qPCR and found to have sizable decreases in percentage of Zika RNA detected when compared with the control (Fig. 4C). Altogether, these results suggest that Zika infection and transmissive capabilities are compromised due to overexpression of CD9, CD63, and CD81.

**Fig. 4.**
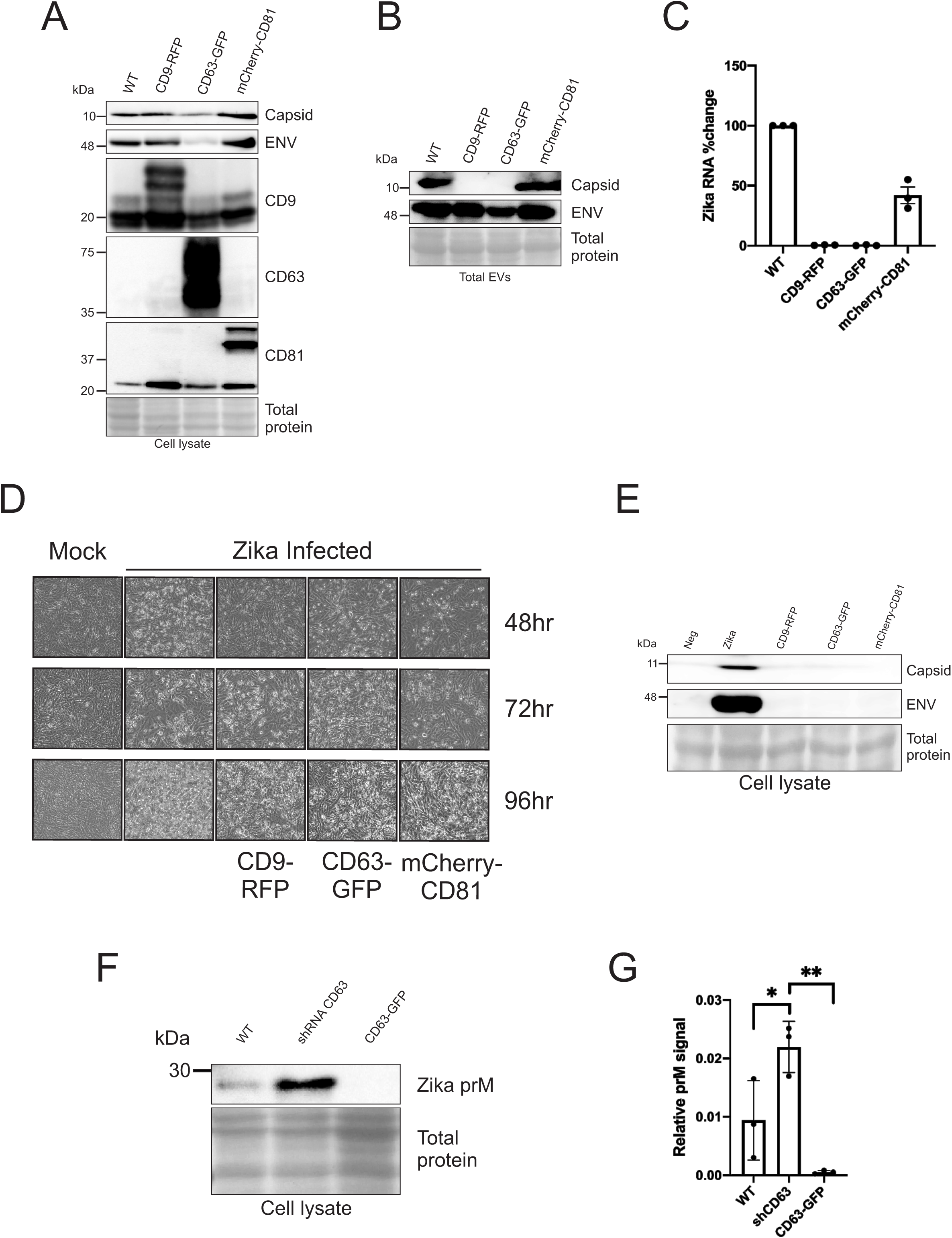
Overexpression of tetraspanin levels decreases Zika infectivity and transmission. A) B) Immunoblots of cell lysate harvested 72 hours from uninfected or Zika infected tetraspanin overexpression and control cells that were probed for tetraspanins and Zika proteins. B) Immunoblot of total EV pellet from infected control and tetraspanin overexpression cells probed for Zika capsid and envelope (total EVs obtained from PEG precipitation of media of uninfected/infected cells following 1000 x g spin for 10 min). C) Quantitation of the percent change in Zika total EV RNA levels of 3 independent experiments (error bars display SEM of 3 independent experiments; samples normalized to control and control set to 100). D)Tetraspanin overexpression and control cells were monitored for CPE via live cell imaging every 24 hours for 4 days post-infection with Zika or mock. E) Immunoblot of cell lysate from cells exposed to EVs from tetraspanin overexpression and control cells. F) Immunoblot of Zika prM protein and G) quantification of 3 independent samples of WT, CD63 knockdown and overexpression cell lysates. Statistical significance was determined using one-way ANOVA (* p<0.05; **p<0.01).

Since tetraspanins appear to regulate vesicle, capsid, and genome release, we further wanted to assess whether knockdown alters infectivity and transmission. The overexpression cell lines were infected with a MOI of 0.5 and CPE was documented by live cell imaging for 4 days (Fig. 4D). CPE was observed but appeared to progress slower in all three cell lines when compared with the control (Fig. 4D). EV transfer experiments were then performed and harvested the cells 96 hours post exposure. Immunoblot analysis of the lysate from the cells exposed to the overexpression tetraspanins EVs all had striking decreases in Zika capsid and envelope proteins (Fig. 4E). These data suggest that EV tetraspanins CD9, CD63, and CD81 modulate Zika replication and spread.

In order to further evaluate how tetraspanins effect replication and/or egress, the levels of Zika prM protein were compared in the CD63 knockdown and overexpression cell lysates (Fig. 4F). We chose to focus on the effects of CD63 since this tetraspainin had the most significant impact on infectivity in the previous experiments. Zika prM protein is the immature form of the structural M protein of the virus particle which is cleaved upon extracellular release. Interestingly, the level of prM is significantly increased in the knockdown and decreased in the overexpression (p<0.05 and p<0.01 respectively) (Fig. 4F-G). These results additionally support that CD63 plays some role in Zika virus replication.

### CD63 localization is altered with Zika virus infection and the expression levels of CD63 effect the cellular localization of Zika capsid protein

Since varying levels of CD63 appear to alter Zika replication and capsid release, we next examined the localization of endogenous CD63 and Zika capsid in the mock and Zika infected cells. Surprisingly, upon Zika infection, CD63 is sequestered to the core of the perinuclearly located viral replication site and surrounded by capsid protein (Fig. 5A). This was a remarkable finding since CD63 was found to be dispersed throughout the cytoplasm of the mock cells (Fig. 5A). These results shed new light on the influence of CD63 on the viral replication site and virally-induced cytoplasmic reorganization.

**Fig. 5.**
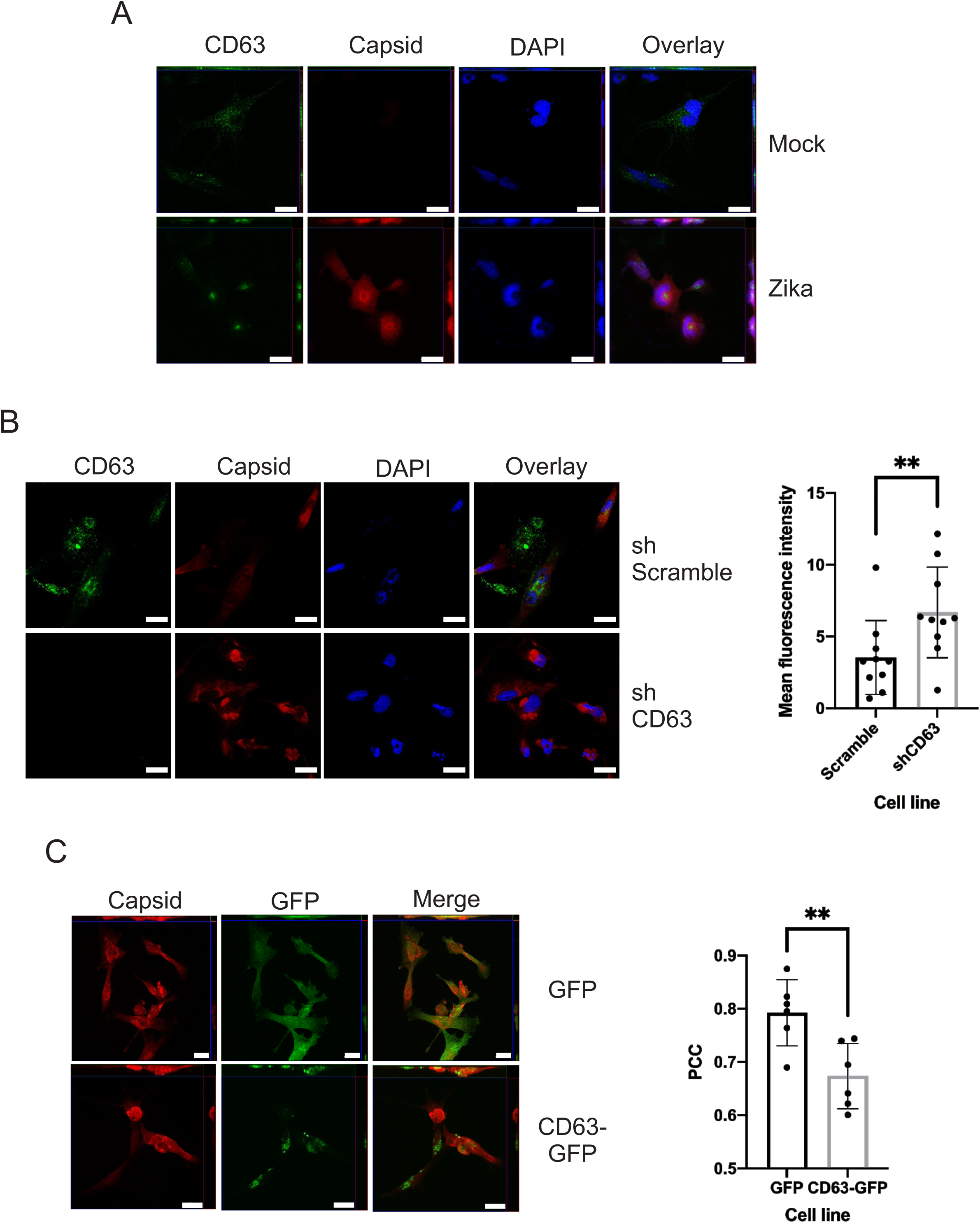
CD63 localization is altered with Zika virus infection and the expression levels of CD63 effect the cellular localization of Zika capsid protein. A) IF of fixed SNB-19 cells was used to analyze endogenous CD63 and Zika capsid localization in WT cells 48 hours mock or post infection with Zika virus. B) SNB-19 inducible shRNA scramble or CD63 knockdown cells were fixed for IF for Zika capsid proteins 48 hours post-infection. C) Representative images and image quantification of Zika capsid localization in GFP control and CD63-GFP overexpression fixed SNB-19 cells 48 hours following infection with Zika virus.

To further ascertain the role of CD63 in Zika infection, Zika envelope and capsid protein localization was assessed in the SNB-19 CD63 knockdown and overexpression cells. When CD63 was knocked-down, capsid was found to form tight cytoplasmic viral replication sites when compared with the scramble control (Fig. 5B). However, in the CD63 overexpression cells, these sites were only observed in cells that lacked or had low CD63-GFP expression (Fig. 5C) In fact, the cells that had high CD63 expression, capsid protein levels were found to be minimal (Fig. 5C). Yet, low levels of CD63 expression revealed a similar phenotypic localization that was observed when endogenous CD63 levels were probed in the WT cells (Fig. 5A,C). Image quantification of the correlation coefficient of GFP/capsid and GFP-CD63/capsid revealed significant decreases in CD63/capsid correlation (Fig. 5C). These data suggest that elevated levels of CD63 disrupt or prevent the formation of capsid containing replication sites. Taken together, these results further support CD63 as having a regulatory role in Zika virus replication and the production of virions and Zika modified EVs.

## DISCUSSION

As the field of EVs continues to expand, there has been accruing interest in the pathological role of EVs in infectious diseases. Numerous studies have recently documented the utilization of EVs in the life cycle of various viruses (16, 17, 57–60). This is not overly surprising since viruses are obligate intracellular parasites and EVs are among the normal processes of cells. Yet, much of the underlying molecular mechanisms behind how viruses utilize these pathways and for what purpose, are still being investigating. We sought to delve deeper into these questions and not only characterize the modification of vesicles by Zika virus but also explore the importance of specific EV trafficking and biogenesis proteins in Zika virus infection and transmission.

Although Zika virus has been recently found to secrete infectious EVs, a detailed systematic analysis of the effects of Zika virus infection on EV-subpopulations has not been performed. Zhou and colleagues released their findings that EVs harvested from Zika infected cells contain Zika genomes and are infectious (61). Our data support these findings but expand on them by demonstrating that Zika infection modifies EV biogenesis and cargo. Furthermore, we show the diversity of vesicle populations released from Zika infected cells, all of which could have distinct functions and influence pathogenic outcomes. Interestingly, van der Grein et al., reported that Picornaviruses, which are also RNA viruses, utilized different EV biogenesis pathways for the secretion of infectious particles with distinct cargo and roles in infection (60). Hence, these discrete vesicle populations may have various roles in maintaining Zika infection and continuing the viral life cycle. Although, alphaviruses have been previously demonstrated to release infectious microvesicles that are lacking capsid protein (62), to our knowledge this is the first report of Zika infected cells releasing infectious small EVs that are capsid-deficient. While we cannot completely rule out the possibility that the small light EVs purified do not contain some virions that produce active infection, virion contamination alone cannot account for the high level of genomes and infectivity found in this EV subpopulation. Other lines of evidence that support a capsid-less EV subpopulation that is infectious include; 1) the inability to detect capsid in this fraction; 2) Zika genomes were detected in this fraction that were sensitive to RNAse treatment following membrane disruption; 3) the inability to detect capsid containing virions by EM; 4) and the ability to pull-down capsid-deficient infectious EV population with PS from the 100k pellet. Regardless of whether capsid-less EVs transmit Zika infection, it is clear from our work and others that Zika virus dramatically modifies EVs produced by cells which likely contributes to Zika spread and pathogenesis within the host.

As mentioned, Zika virus has a complex life cycle with its reliance on vectors as its primary means of transmission. But this virus also has a broad tropism and the capacity to cross cellular barriers such as the blood brain barrier (BBB) and placenta. It is these capabilities that have led to the severe manifestations of the disease including congenital microcephaly. In 2016, Tang el al., demonstrated that Zika virus could infect neural progenitor cells, which leads to cell cycle disfunction and cell death (63). These results reveal how Zika is able to damage the developing fetal brain but the important question of how the virus can cross the placenta and infect the fetus remains unanswered. Interestingly, studies indicate that Zika virus is unable to infect placental trophoblasts and these cells secrete factors that support an antiviral cellular environment within the placenta (64, 65). However, EVs have been reported to cross the placental barrier (23, 24). Here, we have demonstrated that Zika utilizes different biogenesis pathways in cells for the production of virions and virally-modified EVs. Consequently, it is possible that the Zika-modified EVs have different tropic capabilities.

Recent work from Zhou and colleagues revealed that neutral sphingomyelinase 2 (nSMase-2) was required for efficient viral persistence and transmission in mouse cortical neurons (61). Huang and colleagues also showed that treatment with the nSMase-2 inhibitor GW4869, negated the Zika induced EV secretion and reduced the infectivity (66). These findings support EV biogenesis pathways as having a significant role in the transmission of Zika virus since ceramide, the target of nSMase-2, drives budding of vesicles into the MVB (67). Ceramide is implicated in the ESCRT-independent EV biogenesis pathway because it is believed to spontaneously produce budding events when concentrated in the membrane due to the curvature induced from the rigid cone-shaped structure of the lipid head group (67).

Although in this study, we focused primarily on the tetraspanin-mediated trafficking and biogenesis pathways, it is worth noting that it will be important in future studies to examine the role of ESCRT components in Zika infection as tetraspanins can follow ESCRT-dependent and - independent pathways. Other studies have reported the importance of ESCRT proteins in Flavivirus propagation and egress (55, 68). In the study by Thepparit et al., apoptosis linked gene-2-interacting protein X (ALIX) was discovered to be upregulated in cells infected with dengue virus (55). Dengue infection was also shown to be enhanced upon overexpression of ALIX and restricted when ALIX was knocked down in the cells (55). In addition, Tabata and colleagues found that knocking down tumor susceptibility gene 101 (TSG101) and other specific ESCRT-III proteins resulted in significant decreases in dengue and Japanese encephalitis virus titers (42). Therefore, it is quite plausible that there are essential functions for these ESCRT proteins in the Zika virus life cycle. It would be also insightful to explore the role of these proteins in both virion and Zika-modified EV secretion.

In addition to ceramide and ESCRT proteins, tetraspanins have been found to be crucial to other viruses and the secretion of virally-modified EVs (12, 41). For instance, our lab previously demonstrated that CD63 is required for the efficient secretion of Epstein-Barr (EBV) viral oncoprotein latent membrane protein 1 (LMP1) in EVs (12). Tetraspanins have also been implicated in the budding and egress of many viruses including HIV, HCMV, influenza, hepatitis A, HSV-1 and others. In addition, tetraspanins serve important functions in virus entry. For example, hepatitis C virus utilizes CD81 to infect hepatocytes (69). Recently, Vora and colleagues also discovered that dengue infected mosquito cells release tetraspanin (Tsp29Fb) enriched EVs that are infectious to both mosquito and human cells (70). Since the initial characterization of the Zika modified EVs found alterations in tetraspanin protein levels in the EVs from the Zika infected cells, it is possible that these proteins have a role in Zika virus infection and transmission.

The specific contributions of tetraspanins to Zika virus replication can be difficult to decipher due to the complexity of the roles and functions of tetraspanins in cellular processes. Tetraspanins have been shown to form protein and lipid clusters in membranes called tetraspanin enriched microdomains (TEMs), that are thought to serve as platform for EV cargo to congregate as well as sites for virus budding and cell entry (35, 71, 72). It is possible that excessively high or low CD9, CD63, or CD81 levels may disrupt the composition of these TEM sites leading to changes in cargo recruitment to membrane microdomains. This may result in the accumulation of components utilized by the virus thus allowing for more efficient virion production. Alternatively, if a large portion of cellular resources is being funneled through specific EV pathways due to the overexpression of tetraspanins, the virus may be unable to adequately recruit the materials needed for replication and virion assembly.

In addition to virion assembly, it is also possible that the altered membrane composition may affect viral binding, endocytosis or fusion during entry events. The interferon-induced transmembrane proteins (IFTMs) have been reported to accumulate in TEMs as an innate immune response (73). Therefore, it is possible that this accumulation of IFTMs in the TEMs could disrupt tetraspanin interactions and receptor recruitments leading to the restriction of viral entry (74). Furthermore, the availability of cellular receptors or endocytic machinery may be affected by alterations in tetraspanin levels thereby suppressing virus entry.

Our lab previously reported that CD63 has a regulatory role in autophagy and that CD63 bridges the endosomal and autophagic pathways (45). Interestingly, the study by van der Grein and colleagues, found that the Picornavirus light (EV) infectious population was enriched in autophagic markers and indicated that these viruses promote autophagy in cells (60). We just identified that CD63 levels in the different vesicle populations varied when the cells were infected with Zika virus and Zika has also been reported induce autophagy (75). Consequently, it is possible that Zika is utilizing CD63 in the autophagic secretory pathways for the release of autophagically-derived infectious particles. When we compared the Zika genome and protein levels of the small gradient purified EVs from the cells with the overexpression and knockdown of CD63, we found the levels to be significantly less. These finding suggest that specific level of CD63 may be required for the secretion of these Zika-modified EVs.

Interestingly, the immunofluorescence results of endogenous CD63 levels revealed that endogenous CD63 is located in the core of the virus replication unit. Notably, a study by Cortese and colleagues demonstrated that the core of the virus replication unit contains a pore where newly synthesized Zika positive-sense RNA is released (76). When we examined the localization of the Zika envelope and capsid proteins in the CD63 knockdown and overexpression cells, CD63 levels appeared to negatively correlate with capsid levels further supporting high levels of CD63 as being disruptive to the establishment of the Zika virion replication sites. These findings support our hypothesis that CD63 levels may alter the cellular trafficking pathways and assembly sites utilized by the virus for the production and secretion of different types of infectious particles.

It appears that CD63 is a means by which Zika can balance the secretion of virions and infectious EVs. Having lower levels of virion release and infectious Zika EVs may provide better immune evasion and persistence as well as the ability to cross cellular barriers (77). Evolutionary long-term strategies tend to favor adaptability and persistence especially for viruses with complex life cycles such as arboviruses. Arboviruses require a vector to maintain the life cycle but having other routes of transmission, i.e. vertical or sexual, increases the chance of transmission. This would be evolutionarily favorable and Zika virus appears to be exploiting EV biogenesis pathways to increase its transmission and persistence capabilities.

Overall, this study provides a detailed analysis of the extracellular vesicles released from Zika infected cells and new insights into Zika virus cell-to-cell transmission. It also provides the first evidence for the importance of host tetraspanin proteins in Zika replication and spread. Obtaining a better understanding of how Zika virus utilizes tetraspanins and EV pathways to transport infectious genomes and viral and host cargo is likely to offer novel targets to control infection and disease.

## Funding

This study was supported by a grant from the Florida Department of Health (7ZK16) and a Pilot Grant from National Institutes of Health under Award Number U19AI131130. The content is solely the responsibility of the authors and does not necessarily represent the official views of the Florida Department of Health or the National Institutes of Health.

## Authors’ contributions

SBY and DGM conceived of and designed the experiments detailed in this study. MRC, ASC, and LS aided with experimental design. SBY performed the majority of the experiments, with assistance from ASC, LCD and LS. The manuscript was written by SBY and DGM, with equal editing contributions from ASC, LS, LCD and MRC. DGM supervised the project and obtained the funding support. All authors read and approved the final manuscript.

## Acknowledgements

We thank Dingani Nkosi for his help in the construction of the shRNA-expressing plasmids used throughout this study, the Biological Science Imaging Resource at Florida State University for assistance with electron microscopy. The confocal images were obtained with the support of the Florida State University College of Medicine Confocal Microscopy Laboratory and help from Ruth Didier.

